# Characterizing yield through wheat’s perception of chronological progression: a multi-omics plant-time warping approach

**DOI:** 10.1101/2025.05.12.653430

**Authors:** Lukas Roth, Juan M. Herrera, Lilia Levy Häner, Didier Pellet, Dario Fossati, Mike Boss, Xiaoran Chen, Paraskevi Nousi, Michele Volpi

## Abstract

To address challenges in food security, a better understanding of crop performance under varying and changing environmental conditions is required. Plant Time Warping (PTW) is a deep learning model that integrates high-throughput field phenotyping data with genomic and environmental information to predict wheat yield. PTW leverages image time series, genetic markers, and environmental covariates to learn genotype-specific physiological responses to temperature and vapor pressure deficit. Compared to mere genomic prediction models, PTW demonstrates superior performance when predicting yield in unseen environments across 48 year-locations in Europe. The PTW model captures non-linear growth responses varying with phenological stages and identifies distinct patterns associated with yield performance and stability. Specifically, varieties with higher yield stability exhibit reduced sensitivity to vapor pressure deficit around 1.5 kPa and distinctive temperature responses during emergence and senescence. The learned response pattern enable retrospective and prospective yield predictions, providing a foundation for location-specific variety recommendations and targeted breeding strategies. The integration of phenomic, genomic, and enviromic data has the potential to substantially advance research in climate adaptation strategies for crop production by addressing generalization challenges of predictions to novel environmental conditions.

**Highlight:** We present a novel deep learning model that seamlessly combines high-throughput image data, genomic data, and weather data, enabling better crop predictions for future climates.

## 1 Introduction

Three major factors influence crop production systems, and thus impact our global food security: (1) the genetics (*G*) of plants, determined by the choice of variety a breeder selects and farmer grows, (2) the environment (*E*) that the growing plants experience, and (3) the management practices (*M* ) applied by the farmer. In the breeding and variety testing context we address here, *M* is generally considered to be optimized beforehand and hence a constant, leaving us with *G* and *E* as model inputs to understand the dynamics of such systems. Considering potential interactions between these factors, a common starting point is to define a phenotype *P* to be the result of genotype effects *G* with complex environments *E* (Malosetti et al. 2016),

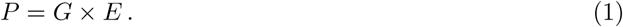

Describing *G* with molecular methods is nowadays feasible down to the whole-genome level (Walkowiak et al. 2020). Public weather station data to describe *E* are becoming increasingly available via governmental institutions that commit themselves to open data (e.g., Deutscher Wetterdienst (DWD) and MeteoSwiss). In contrast, public variety testing data and other multi-environment trial (MET) data to describing *P* s at a set of *E*s are yet rare.

However, while this lack of data is a restriction, it is not the main cause for the current limitations of predictive models. The key is that *G* and *E* are measured at massively different scales than *P* . While data sets comprising genetic marker data and time series of environmental covariates easily exceed 100k data points (Roth et al. 2024b), MET data barely reach the 10k level (Herrera et al. 2020). Taking the Swiss winter wheat variety testing as an example, the ten locations and 36 examined varieties per year would only add up to 3,600 data points if 10 years of data were used. The result is a typical ‘large *p*, small *n*’ problem, where one tries to model system dynamics with many variables (*p*) but few samples (*n*).

Field phenotyping technologies promise to address this problem by time-resolving *P* in *P_t_* measurements that describe the phenotype *P* at time *t* (Roth et al. 2021). Hence, one can define *P_t_* as time-integrated result of genotype effects *G*—assuming fixed gene expression over the growing season—with complex and changing environments *E_t_*,

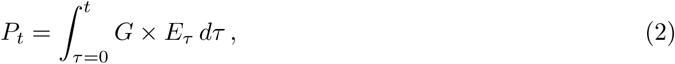

where *τ* is time over the growing season and *t* the measurement point in time. Field phenotyping can deliver data sets with 150k data points and more (Roth et al. 2024b), which brings data set sizes of *P_t_* to the level of *G* and *E_t_*, allowing to find empirical solutions for Equation 2.

The question arises how to model the interactions between *G* and *E*. Historically, such *G × E* effects are considered to be the product (*×*) of scalars describing *E*s (e.g., mean yield performance per *E*) and scalars describing *G*s (e.g., factors of a Finlay-Wilkinson regression) (Fisher and Mackenzie 1923). Yet, it has been shown that genotype effects can be further detailed in small effects of deoxyribonucleic acid (DNA) polymorphism (Li et al. 2021) that can be integrated in whole-genome predictions approaches (Meuwissen, Hayes, and Goddard 2001). Similarly, environment effects can be further detailed using environmental covariates such as temperature and precipitation in whole-genome prediction approaches (Jarquín et al. 2014).

Small effects of *G* and *E* interactions are then additively combined to predict the performance of a specific genotype in a specific environment. Hence, a linear relationship between genetic and environment characteristics is implied. Yet, strong evidence point towards the fact that relations between environmental covariates and growth are far from being linear (Roth et al. 2024a). Additionally, these relationships change over time as crop phenology advances (Porter and Gawith 1999). Since crop growth and phenological development are primary drivers of overall crop performance, the assumption of constant linearity appears to be an oversimplification (Wang et al. 2017).

Consequently, it was suggested to combine whole-genome prediction approaches with crop growth models that model non-linear dependencies based on *a priori* knowledge (Diepenbrock et al. 2021). However, such approaches rely heavily on pre-calibrated model parameters (van Voorn et al. 2023). When crop models are used in a breeding context, their genotype-specific parameters are typically reduced to a handful, while the rest remain fixed (Chapman 2008). Moreover, crop models may even require additional controlled experiments to further refine predictions to a specific set of environments (Millet et al. 2019). In this work, we hypothesize that the bias of linear models and the out-of-domain issues of pre-calibrated crop growth models can be avoided by directly learning crop growth models on time series of phenotyping data, that is on *P_t_* data instead of *P* data.

Attempting to integrate *P_t_* values in any model raises the question of how to represent growth. While phenotyping data are usually multidimensional, e.g., imaging data or 3-D point clouds, common modelling approaches integrate growth as a scalar, e.g., a growth rate (Roth, Piepho, and Hund 2022; Roth et al. 2024a). Hence, a common step is to first apply feature extraction and dimensionality reduction techniques to multi-dimensional phenotyping data (Hund et al. 2019). Often, such approaches follow a domain-specific purpose, i.e., they try to approximate a physical trait such as for example number of plant organs, canopy structure, leaf area, and others. However, choosing which traits to track over time may create a strong bias towards identifying only well-established relationships.

To address these issues, we propose a novel end-to-end deep learning model called plant time warping (PTW) that is trained simultaneously on high-resolution red, green, and blue (RGB) images of hundreds of wheat genotypes grown in the field phenotyping facility ‘FIP’ of ETH Zurich, and on genetic marker data and environmental covariates (Figure 1). The PTW model inherently learns (1) a representation of growth from images (Figure 1, Loss 1 and P-state), and (2) how to represent interactions between genetic marker and environmental covariates that explain this growth (Figure 1, Loss 2/3 and G*×*E-state).

**Figure 1:**
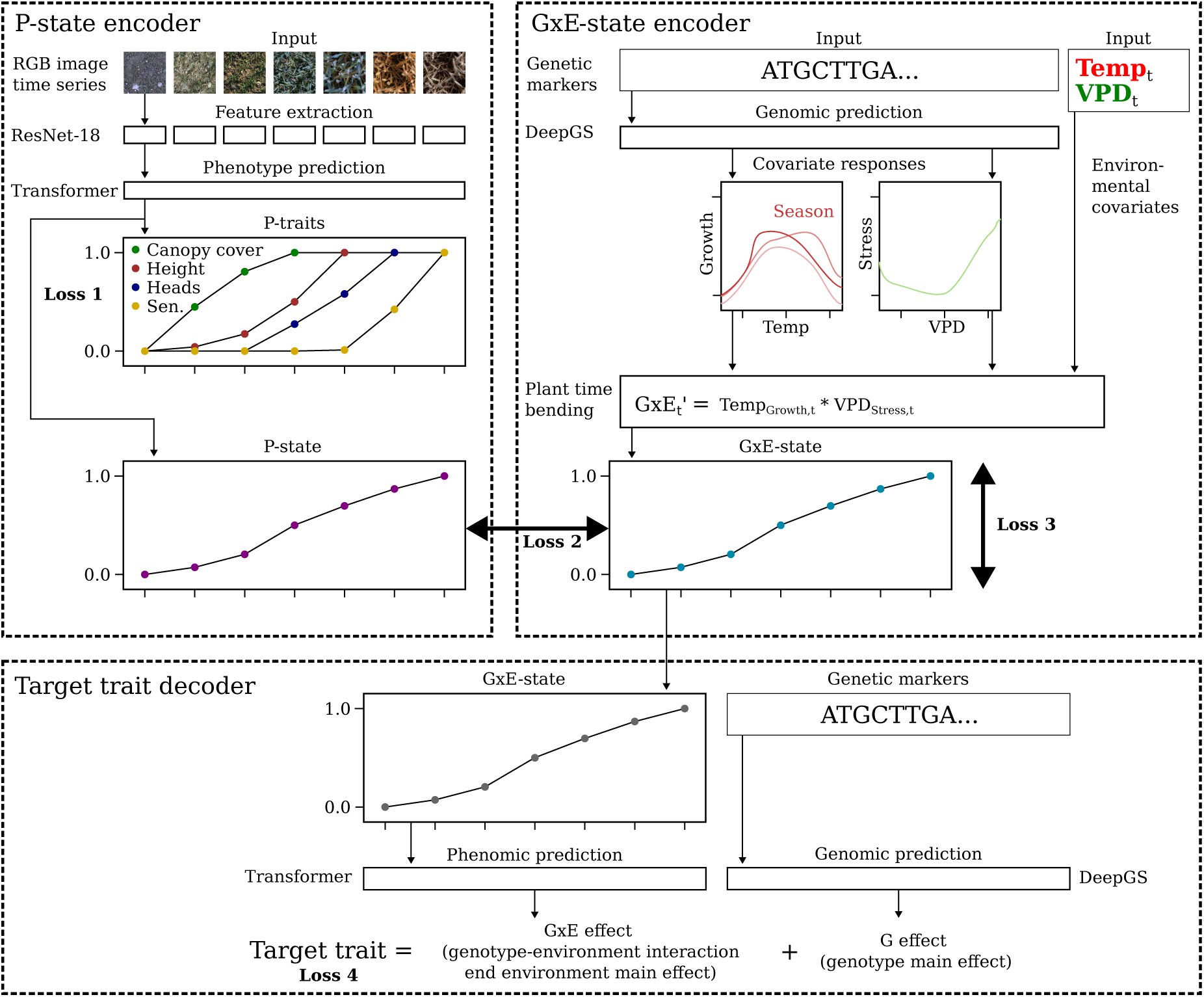
Plant time warping (PTW) model with P-state encoder, G*×*E-state encoder, and target trait decoder. Input to the P-state encoder are image time series, and to the G*×*E-state encoder environmental covariates (temperature, vapour pressure deficit (VPD)) and genetic marker data as single-nucleotide polymorphisms (SNPs). the similarity of two temporal sequences that vary in speed, i.e., the cost of ‘warping’ them to each other. Here, we consider non-linear plant growth within a growing season as a ‘warped’ representation of cardinal time. Unlike homeothermic species such as humans, plant growth strongly depends on environmental covariates. Hence, with our approach, we intend to learn a genotype-specific warping function of cardinal time to ‘plant time’ in dependence of environmental covariates, and consequently, we name this method ‘plant time warping’.

The PTW model is built around the philosophy of ‘warping’ time with environmental covariates in a way it represents non-linear genotype-specific growth (Roth et al. 2024a), and subsequently using this learned latent representation of growth to predict yield. The term ‘time warping’ was shaped in the context of pattern recognition for time series (Berndt and Clifford 1994), where one wished to describe

Based on unseen data from 48 year-locations in France, Germany and Switzerland, we validate the PTW model and test its ability to predict unseen environments, unseen genotypes, and unseen genotypes in unseen environments. Furthermore, we evaluate the model on 20 years of historic weather data and yield measurements on a country scale. Finally, we use the trained model to extract optimized phenotypes, so-called ideotypes, that promise higher and more stable wheat yields.

## 2 Materials and methods

### 2.1 Data

Three different data sets were used to train and test the PTW model, (1) the FIP 1.0 data set (Roth et al. 2024b), (2) the GABI-MET data set (Gogna et al. 2022), and (3) the CH-MET data set (Herrera et al. 2020). Details of the three data sets are described in the following. Please note that, although all three data sets contain plot-specific measurement data, environmental covariates are measured at a weather station near the field site, in line with current best practice in field phenotyping.

#### 2.1.1 FIP 1.0 data set

The FIP 1.0 data set consisted of annotated image time series acquired with the Field Imaging Platform (FIP) at ETH Zurich in six years (Roth et al. 2024b). The open data set comprises approximately 4,000 image time series of wheat plots. Each image in the time series is capturing a canopy area of 1.0*×*1.5 m of a specific winter wheat genotype. Time series are annotated with the secondary/low-level traits canopy cover, plant height, wheat head counts, and senescence ratings, and with the target traits yield, protein content, heading date, and final height. Of the four low-level and four target traits, all low-level traits but only the target trait yield were used for the PTW model.

Also part of the FIP 1.0 data set are genetic marker data (90k SNP array) for the GABI-WHEAT panel based on the public data set published by Gogna et al. (2022). A separate set of additional marker data for genotypes that are not part of the GABI-WHEAT panel was generated using a 45k SNP array (non-public data). Hence, two sub-data sets were assembled: one consisting solely of GABI-WHEAT panel markers and another extended set that combined overlapping markers from both public and protected sources. Marker data filtering and imputation procedures are part of the data set preparation done by Roth et al. (2024b). The final GABI-WHEAT panel marker data set after filtering comprised 372 genotypes (GABI-WHEAT panel) and 18,846 markers. The extended data set after filtering consisted of 844 genotypes (GABI-WHEAT panel, registered Swiss varieties, and F8 generation genotypes) and 11,943 markers.

Image time series in the FIP 1.0 data set are annotated with environmental covariates from a local weather station. For the PTW model, air temperature and relative humidity were used, while vapour pressure deficit (VPD) was derived from temperature and relative humidity assuming an atmospheric pressure of 101 Pa.

#### 2.1.2 GABI-MET data set

The GABI-MET data set consists of plot-based target traits and derived adjusted genotype values (best linear unbiased estimations (BLUEs)) for eight environments as part of a multi-environment trial (MET), and genetic marker data for the GABI-WHEAT panel (see previous section) (Gogna et al. 2022). BLUEs of MET data from the eight year-locations were used as first test set for unseen environments. The MET data were enriched with hourly temperature and relative humidity data originating from the Climate Data Center (CDC) of the German Weather Service (for German environments), and from Meteo France (SYNOP, 3-hourly data only) (for French environments). The data set comprised 300 genotypes that were part of the FIP 1.0 data set and therefore seen in training, and 60 other genotypes that were unseen in training.

#### 2.1.3 CH-MET data set

MET data from 40 year-locations (years 2010–2013, 10 locations) from the official Swiss variety testing (Agroscope, Switzerland) were used as second test set for unseen environments (Herrera et al. 2020). Experimental field designs followed a Latin square design with three replications and 36 genotypes per year-site (only 24 genotypes in 2010). Genotype overlaps between years were in minimum 15% (2010 to 2013) but otherwise between 25 and 50%. BLUEs were calculated based on plot-based values using ASReml-R (The VSNi Team 2023) with genotype and replicate as random factors and row and columns

The CH-MET data included 71 genotypes in total. For 35 genotypes, genetic marker data were available, 22 of these genotypes were also part of the FIP 1.0 data set and hence seen in training. The remaining 13 genotypes were unseen in training.

### 2.2 Model architecture

The PTW model consists of two encoder parts (P-state and G*×*E-state encoder) and one decoder part (target trait decoder) (Figure 1). The P-state encoder takes image time series as input and predicts time series of P-traits (canopy cover, plant height, wheat head counts, and senescence) and time series of P-states (latent representation of growth of the measured phenotype). The G*×*E-state encoder takes genetic marker data (SNPs) and time series of environmental covariates (temperature and VPD) as input and predicts time series of G*×*E-states (latent representation of growth of the modelled phenotype). The P-state from the P-state encoder corresponds to the G*×*E-state from the G*×*E-state encoder in regard to representing the same characteristic (growth) with different means (phenotyping data versus genomic and environmental data). The target trait decoder takes P-state/G*×*E-state time series and genetic marker data and predicts yield. The two encoders are trained in parallel to agree on the P-state/G*×*E-state, following a concept inspired by recent trends in self-supervised, non-contrastive learning from computer vision and large language model research (Tian, Chen, and Ganguli 2021).

The PTW model was implemented in pytorch (Paszke et al. 2017) and trained with pytorch-lightning (Falcon and team 2025). Common to all model parts are rectified linear unit (ReLU) activation functions (Glorot, Bordes, and Bengio 2011), a common choice for transformer-based architectures. Further details on specific model parts are described in the following.

#### 2.2.1 P-state encoder

Feature embedding of images per point in time was achieved using a ResNet18 architecture, with weights pretrained on ImageNet (He et al. 2015), where the last classification layer was removed, yielding a 512-dimensional output vector. This representation served as an initial input for a transformer model.

ResNet18 was chosen because it is a lightweight, proven architecture to compress high-dimensional input into a compact feature vector with high robustness.

The transformer model (Vaswani et al. 2023) consisted of 32 attention heads and 16 layers. Positional encoding was added to the input embeddings using sinusoidal functions as proposed in Vaswani et al. (2023). Transformers provide attention across time if working with time series while respecting temporal order if positional encoding is provided.

Subsequent to the transformer, four fully connected layers were applied with dimensions 512, 128, 6, and 5. Note that the second-last layer has only one extra dimension more than the output layer (6 versus 5). By forcing this low dimensionality, we intend this layer to learn a statistical representation of growth patterns related to all low-level traits.

For the output layer, a sigmoid activation function was used to normalize trait predictions (canopy cover, plant height, head counts, senescence, and P-state) into [0, 1]. While the four traits were trained in a supervised way using labels, the fifth output value, the P-state, was intended to represent an overall representation of growth, i.e., a statistical representation that facilitates the prediction of yield. This trait was trained in a self-supervised manner by simultaneously predicting its correspondence, the G*×*E-state, with the G*×*E-state encoder based on environmental covariates and genetic marker data, as described in the following.

#### 2.2.2 G*×*E-state encoder

Time series with hourly measurements of temperature and VPD and genetic marker data were input of this second encoder. The choice to model growth as an immediate response to temperature and VPD was based on two major considerations. First, research on modeling growth responses to environmental covariates using temporally highly resolved time series has identified temperature and VPD as important drivers (Walter, Silk, and Schurr 2009). These findings from indoor platforms were recently confirmed for the Swiss environment using field-based data (Roth et al. 2024a). Second, following the principle of parsimony in crop modeling (Hammer et al. 2019) requires keeping the number of covariates low. Temperature and solar radiation, as well as precipitation and VPD, are highly correlated covariates. Focusing on widely available temperature and VPD measurements ensures transferability of the model architecture to diverse data set scenarios.

As encoder for the genetic data based on SNPs, the DeepGS model architecture suggested by Ma et al. (2018) was used. DeepGS is a deep learning genomic prediction model based on a lightweight standard convolutional architecture that has shown comparable performance to state-of-the-art genomic prediction approaches such as GBLUP.

Two separate DeepGS model parts were included in the encoder to estimate temperature growth response parameters and VPD stress response parameters. The DeepGS architecture followed the original publication, with an additional batch normalization layer before the first layer.

A first set of spline knot coordinates predicted by DeepGS was used to parametrize a 2D spline fitting growth response to temperature and days after sowing, i.e., changing responses to temperature with time. A second set of spline knot coordinates predicted by DeepGS was used to parametrize a 1D spline that fitted stress response to VPD, i.e., a constant penalization of growth with VPD.

The G*×*E-state *G×E_t_*at point in time *t* was computed as a cumulated response to temperature and VPD,

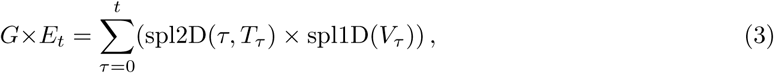

where spl2D represents a 2D spline with 10*×*8 knots and spl1D a 1D spline with 8 knots, *V* denotes VPD, and *T* denotes temperature.

#### 2.2.3 Target trait decoder

The estimated G*×*E-state values served as input for a transformer-based decoder trained to predict yield. Daily G*×*E-state values were retained up to 260 days after sowing as input tokens, with subsequent values discarded. An additional token was introduced for yield prediction. Positional encoding was applied to the sequence to annotate DAS.

A transformer model was employed with an embedding dimension of 128, 64 attention heads, and 16 layers. The output from the transformer was processed through three fully connected layers (dimensionality 128, 64, 64), yielding the final yield prediction based on G*×*E-state time series.

For the genetic main effect, one additional DeepGS model was included. The final yield prediction was obtained as the sum of the G*×*E effect and the genetic main effect,

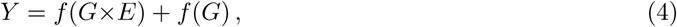

where *f* (*G×E*) is the transformer taking *G×E* time series as input and *f* (*G*) is the DeepGS model taking marker data as input.

### 2.3 Training procedure

The PTW model was trained using a weighted combination of four loss terms (Figure 1), two for the supervised training part (*L*_1_, *L*_4_) and two for the self-supervised part (*L*_2_, *L*_3_),

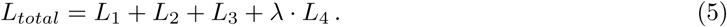

*L*_1_ minimized the mean squared error (MSE) between predicted P-traits (*y_ip_*) and labelled P-traits (*y_ip_*) per trait *p* (canopy cover, plant height, wheat heads, and senescence) and image *i* in image time series,

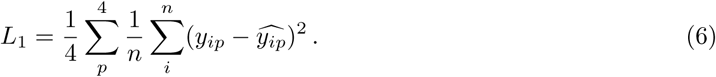

*L*_2_ minimized the MSE between predicted G*×*E-state values (*G ×E*) and predicted P-state values (*P*^-^),

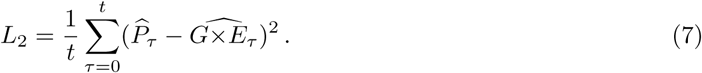

*L*_3_ maximized the range of the G*×*E-state per time series,

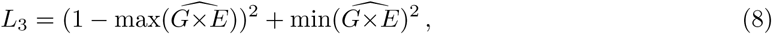

where *G×E* are G*×*E-state values in the range [0, 1]. This loss is required to prevent the PTW model from minimizing *L*_2_ by predicting all-zero for P-state and G*×*E-state, i.e., it enforces

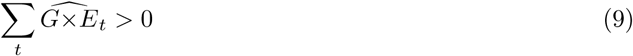

to prevent model collapse. *L*_4_ minimized the MSE between predicted yield values (*y*^) and labelled yield values (*y*) per image time series *s*,

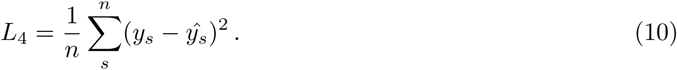

A scaling factor of *λ* = 10 for *L*_4_ was empirically determined to balance its contribution relative to the other loss components, ensuring that the model focuses on accurate yield predictions while no single term would dominate the optimization process.

Training was conducted in two phases. In a first training phase for 16 epochs, the genetic main effect was set to zero, and the PTW trained using all available samples. This phase can be considered as a warm-up phase for the P-state encoder. In a second, significantly longer training phase of 28 epochs, genetic main effects were included while freezing the P-state prediction component. Only samples with available genetic marker data were used in this second phase.

If MET data were included in the training procedure (see next Section), an additional third training phase followed. In this phase, the additional MET data were used to fine-tune the model for another 125 (GABI-MET) respectively 250 (CH-MET) epochs together with the image data.

In contrast to the FIP 1.0 data, the MET data could not be augmented, as they do not contain images. Consequently, only two (for GABI-MET) respectively one (for CH-MET) augmentation per image data sample were used. This ensured a close-to 1:1 ratio between image and MET data.

#### 2.3.1 Training variants

PTW models were trained on either the GABI-WHEAT SNP set (noted as context ‘GABI’) or the extended SNP set (noted as context ‘Extended’). Three training and test scenarios were chosen. PTW models were either trained on (1) the pre-defined training/validation/test split of the FIP 1.0 data set (5 years, 1 location), (2) the full FIP 1.0 data set as training (6 years, 1 location), or (3) the full FIP 1.0 data set combined with either part of the GABI-MET data set or CH-MET data set (7 years, 4 locations, respectively 9 years, 11 locations) as training (Figure 3). Training variants (1) and (2) were independently trained from scratch. Training variants (3) that included GABI-MET or CH-MET data in training were fine-tuned on variant (2) models.

If the full FIP 1.0 data set was used, tests were only performed on unseen MET data. If GABI-MET was additionally included in the training, one year (2009) was included in the training while the other year (2010) served as test set. If CH-MET was additionally included, three years (2010–2012) were included in the training while the left-over year (2013) served as test set.

In six ablation studies, critical components of the PTW model where switched of to show their relative contribution and importance. Those studies included all four losses (L1–L4), the encoder transformer architecture, and the genetic main effect component based on a second DeepGS model. Ablation studies were trained on the FIP 1.0 GABI-WHEAT SNP set data set and compared with the corresponding fully-fledged PTW model.

### 2.4 Data augmentation

Colour and canopy structure variation due to external factors such as wind, moisture, sun orientation and cloud cover is very common in field-based phenotyping data. The following measures were taken to increase the robustness of the models against such noise. Images captured with the field phenotyping platform (FIP) were subjected to a squared random cutout process, extracting 35% *±* 10% of the original image, resulting in a cropped area of approximately 1.3*×*0.875 m (Figure 2). The cutout position was randomly determined within the original image. Additionally, images were randomly rotated and resampled to 224*×*224 pixels. All random operations followed a uniform distribution. The cutout bounding box was kept constant within the image time series, resulting in image patches that showed approximately the same three plant rows throughout the season.

**Figure 2:**
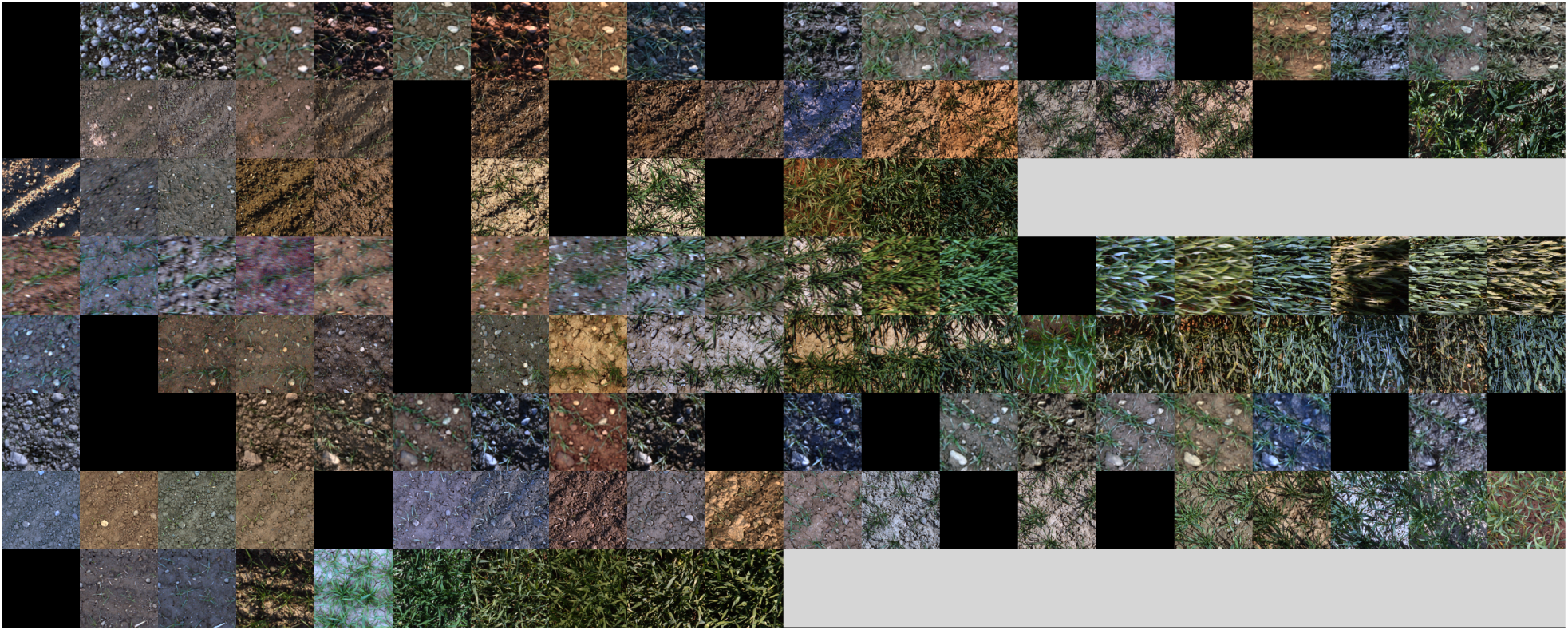
Image input batch example, showing random cutout per image time series (rows), varying time series length (grey images), random masking (black images), and color jittering. Time series are cropped to length 20 to improve visual representation in this manuscript, original time series lengths are <=63.as auto-regression of order 1 (AR1)*×*AR1 residual model to account for gradients on fields. Hourly temperature and relative humidity originated from the MeteoSwiss data base interface Idaweb.

**Figure 3:**
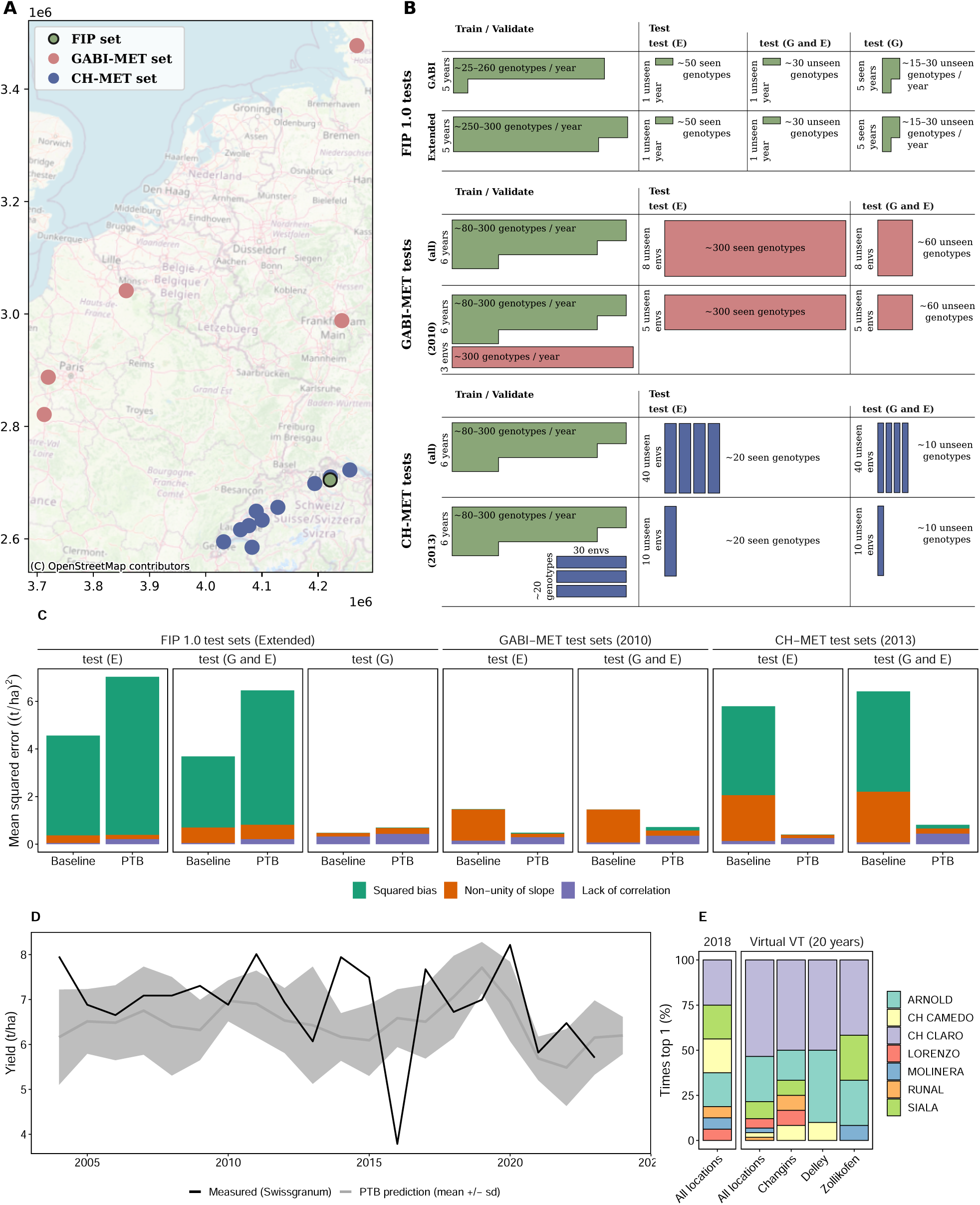
Training/test scenarios (A, B), test performance (C) for the three data sets field phenotyping platform 1.0 (FIP 1.0), GABI multi-environment trial (GABI-MET), and Swiss variety testing multi-environment trial (CH-MET), and 20-years predictions (D, E) of the proposed plant time warping (PTW) model. Test performance for the PTW model and baseline model (random regression) is reported as mean squared error (MSE) decomposed in its three model-prediction related components squared bias, non-unity of slope, and lack of correlation. 20-year predictions show average variety performance 1a1nd standard deviation (sd) (D) and recommended varieties per location (E).

We also applied physics based color augmentation using Kornia’s RandomPlanckianJitter that sim-ulates illumination changes between scenes (probability of 0.8, in “blackbody” mode, selecting color temperatures from the range [4, 15]). Temporal masking of images within the time series was performed at a frequency of 0.1. Each sample underwent 12 augmentations, including the cutout and random color jittering.

### 2.5 Baseline

As baseline served a genomic prediction model based on a random regression to environmental covariates (Jarquín et al. 2014). The FIP 1.0 data set publications provides baselines for the proposed train/validation/test splits. For the additional test sets GABI-MET and CH-MET, identical models as in the FIP 1.0 paper were trained on the full FIP 1.0 data set and on combined FIP 1.0 and MET training sets.

Environmental covariates were the standardized precipitation and evapotranspiration index (SPEI) (Beguería et al. 2014), VPD, air temperature at 2.0 m above ground, and precipitation. For the SPEI and temperature, the mean, maximum and minimum value over the season were used, for VPD, the mean and maximum, and for precipitation, the sum of the values was used. Genomic prediction models were fitted with ASReml-R (The VSNi Team 2023) as described in Roth et al. (2025).

### 2.6 Evaluation metrics

As test metric, mean squared error (MSE), Pearson’s r (correlation) and mean absolute percentage error (MAPE) were used. MSE was decomposed in the three components squared bias, non-unity of slope, and lack of correlation as proposed by Gauch, Hwang, and Fick (2003). Pearson’s r (correlation) and mean absolute percentage error (MAPE) were calculated per environment and then averaged (‘Within E’), and across all environments simultaneously (‘Across E’).

P-state traits were predicted on a plot base, i.e., with replications of genotypes. This allowed to calculate heritability according to Oakey et al. (2006) with SpATS (Rodríguez-Álvarez et al. 2018).

### 2.7 Model analysis and interpretation

#### 2.7.1 Ideotype clustering

To visualize optimized phenotypes in regard to yield (so-called ideotypes), the learned responses to temperature and VPD of the PTW model trained on the full FIP 1.0 data set were clustered according to their shape. K-shape clustering (Paparrizos and Gravano 2016) with cluster sizes *k* = [4, 5, 6*, …,* 9, 10, 20*, …,* 80] were evaluated regarding their Dunn index (Arbelaitz et al. 2013) with the R package dtwclust (Sarda-Espinosa 2023). Clusters built by the winning configuration (*k* = 40) were annotated with yield performance and yield sensitivity mean values evaluated on the GABI-MET data set with a Finlay-Wilkinson regression (Finlay and Wilkinson 1963) performed with the R package statgenGxE (van Rossum 2023). Three clusters that were at the extremes of the yield performance versus yield sensitivity space were visually identified. The targeted ideotype, a cluster with high yield and low yield sensitivity, was contrasted to two other clusters, one with low yield and low yield sensitivity and one with high yield but high yield sensitivity.

#### 2.7.2 Yield prediction and variety recommendations for 20 years

To demonstrate the potential of the learned PTW model to predict for unseen years, genotype performances were predicted for 20 years of environmental data (temperature and relative humidity) from MeteoSwiss. Predictions were performed for the 10 locations of the CH-MET variety testing data set for 14 varieties (quality class TOP and I) that were on the recommended variety list for 2018 (Courvoisier et al. 2017). Variety predictions were then ranked per year for number of times being the top 1 variety per location. Yield per location and variety were averaged to mean yield and variance of yield per year, and compared with published annual yield per area data by swiss granum https://www.swissgranum.ch/zahlen/inlandproduktion. As Swiss granum data and variety testing data are characterized by the typical shift in absolute values between experimentally determined yield and yield measured on true agricultural fields, swiss granum data were aligned to variety testing data by correcting for a fixed offset (−0.63) and slope (2.26) determined in 2010–2014 where data of both types were available.

## 3 Results

### 3.1 Model performance

Common yield prediction models either utilize environmental covariates alongside genomic marker data that characterizes the genetic set-up of crop varieties (genomic prediction) or phenomic data that describes crop phenotypes (phenomic prediction). The proposed PTW model offers a hybrid approach that can be trained on both genomic and phenomic data, yet can generate predictions using genomic data alone. Hence, we trained the PTW model on both phenotyping data (FIP 1.0 data set, MET data set) and genomic data (SNP data set) in combination with environmental data. The PTW model was then tested on previously unseen MET data using only environmental and genomic data as inputs. This testing approach most closely resembles a genomic prediction framework and was therefore compared to a common genomic prediction approach.

Random regression models represent an established approach in genomic prediction that estimates genetic effects as continuous functions across environmental gradients (Jarquin et al. 2018). Such models can handle correlated environmental covariates through appropriate covariance structures (Tolhurst et al. 2022). We benchmarked our novel deep learning architecture against this conventional method.

The genomic prediction model performed well when tested on unseen genotypes, with a MSE of less than 1 t/ha (Figure 3C), corresponding to a MAPE of less than 10 % (Table 1). However, error sizes increased substantially when the test environment was unseen, with a MAPE of approximately 30 % for the GABI-MET data set and around 20 % for the FIP 1.0 and CH-MET data set. Correlations within environments for the MET data sets were weak to moderate (0.35–0.53), for the FIP 1.0 data set very weak to strong (0.15–0.69).

**Table 1:** Performance of suggested plant time warping (PTW) model and of baseline (random regression to environmental covariates) on independent test sets. Provided are Pearson’s r (Cor) and mean absolute percentage error (MAPE) within environments (Within E) and across environments (Across E) for seen genotypes in unseen environments (E), unseen genotypes in unseen environments (G and E), and unseen genotypes in seen environments (G). Models were trained on either the field phenotyping platform 1.0 (FIP 1.0) data set train split, the full FIP 1.0 data set, or the FIP 1.0 data set plus one of the multi-environment trial (MET) data set, GABI multi-environment trial (GABI-MET) or Swiss variety testing multi-environment trial (CH-MET).

| Context | Trained on | Model | Pearson's r (correlation) |  |  | MAPE |  |  |
| --- | --- | --- | --- | --- | --- | --- | --- | --- |
| *FIP 1.0 test sets, GABI SNP set* |  |  | (E) | (G and E) | (G) | (E) | (G and E) | (G) |
| Within E | FIP (train) | Baseline | **0.69** | **0.15** | **0.45** | **24.5** | **19.4** | **8.2** |
|  |  | PTW | 0.32 | -0.08 | 0.35 | 31.5 | 24.8 | 9.4 |
| Across E | FIP (train) | Baseline | - | - | **0.80** | - | - | - |
|  |  | PTW | - | - | 0.68 | - | - | - |
| *FIP 1.0 test sets, extended SNP set* |  |  | (E) | (G and E) | (G) | (E) | (G and E) | (G) |
| Within E | FIP (train) | Baseline | **0.68** | **0.23** | **0.48** | **22.9** | **19.7** | **7.7** |
|  |  | PTW | 0.59 | 0.17 | 0.44 | 29.1 | 27.2 | 8.1 |
| Across E | FIP (train) | Baseline | - | - | **0.80** | - | - | - |
|  |  | PTW | - | - | 0.71 | - | - | - |
| *GABI-MET test sets, extended SNP set* |  |  | (E) | (G and E) | (G) | (E) | (G and E) | (G) |
| Within E | FIP (all) | Baseline | **0.35** | **0.37** | - | 28.4 | 29.2 | - |
|  |  | PTW | 0.15 | 0.01 | - | **20.3** | **22.1** | - |
|  | FIP & CH-MET | PTW | 0.23 | 0.13 | - | 26.1 | 27.7 | - |
| Within E (2010) | FIP & GABI-MET | Baseline | **0.53** | **0.53** | - | 10.9 | 10.9 | - |
|  |  | PTW | 0.41 | 0.25 | - | **5.9** | **7.0** | - |
| Across E | FIP (all) | Baseline | **0.20** | **0.24** | - | 28.4 | 29.2 | - |
|  |  | PTW | 0.10 | 0.02 | - | **20.3** | **22.1** | - |
|  | FIP & CH-MET | PTW | 0.08 | -0.03 | - | 26.1 | 27.7 | - |
| Across E (2010) | FIP & GABI-MET | Baseline | 0.27 | 0.30 | - | 10.9 | 10.9 | - |
|  |  | PTW | **0.85** | **0.80** | - | **5.9** | **7.0** | - |
| *CH-MET test sets, extended SNP set* |  |  | (E / G and E) |  | (G) | (E / G and E) |  | (G) |
| Within E | FIP (all) | Baseline | **0.51** |  | - | **19.9** |  | - |
|  |  | PTW | 0.36 |  | - | 21.5 |  | - |
|  | FIP & GABI-MET | PTW | 0.33 |  | - | 27.7 |  | - |
| Within E (2013) | FIP & CH-MET | Baseline | **0.50** |  | - | 43.7 |  | - |
|  |  | PTW | 0.45 |  | - | **11.2** |  | - |
| Across E | FIP (all) | Baseline | 0.14 |  | - | **20.5** |  | - |
|  |  | PTW | **0.21** |  | - | 23.8 |  | - |
|  | FIP & GABI-MET | PTW | 0.09 |  | - | 27.8 |  | - |
| Across E (2013) | FIP & CH-MET | Baseline | 0.16 |  | - | 43.7 |  | - |
|  |  | PTW | **0.87** |  | - | **11.2** |  | - |

The suggested PTW model consistently outperformed the random regression baseline for the 48 year-locations of the MET data sets. Its performance was superior for both test scenarios, (1) unseen environments (test (E)) and (2) unseen genotypes in unseen environments (Figure 3C, test (G and E)). While the baseline struggled with the larger CH-MET data set with 40 year-locations, exhibiting significantly higher mean squared errors (MSEs) compared to the smaller GABI-MET data set with 8 year-locations, the PTW model effectively reduced MSEs for both data sets to similarly low levels.

The MSE can be influenced by different factors whose relative importance may vary among models. Gauch et al.’s MSE decomposition method provides a valuable framework for evaluating model performance by separating the mean squared error into three distinct components: squared bias (systematic error), non-unity slope (proportional deviation), and lack of correlation (random error) (Gauch, Hwang, and Fick 2003). Although the PTW model tended to increase the lack of correlation on test sets compared to the baseline, this drawback was outweighed by a reduced non-unity of slope and lower squared bias, leading to overall improved performance (Figure 3 and Extended Data Figure 5–7).

While the PTW model was superior for the MET data sets, it underperformed for the FIP 1.0 data set. Higher bias and a slight increase in the lack of correlation contributed to an overall higher MSE than for the baseline across both test scenarios. Pearson’s correlation coefficients and mean absolute percentage errors (MAPEs) confirmed the baseline’s advantage, though the differences were relatively small (Table 1). For instance, in the model using the extended SNP marker set, correlation differences were less than 0.09. Notably, differences between the two SNP marker sets were negligible.

The ablation studies revealed the importance of loss L2–L4 and the genetic main effect in preventing representation collapse—prediction accuracies dropped to close-to zero if switched off. Interestingly, switching off loss L1 increased the accuracy on the FIP 1.0 data set, but also increased the MAPE for the unseen test set (unseen G in unseen E) by 12%, indicating the importance of low-level trait representation in the latent space for extrapolation to unseen conditions. Removing the P-state transformer reduced the MAPE but resulted in deteriorated accuracy for the unseen test set as well.

When evaluating within-environment correlations and MAPE for the MET data sets, the baseline demonstrated slightly better correlations than the PTW model. However, this gap narrowed when models were trained on combined FIP 1.0 and MET data sets. For the MAPE, the PTW model even surpassed the baseline when trained on combined data sets. When evaluation MAPE and correlations across environments, the PTW model was clearly superior to the baseline, provided again it was trained on combined data sets.

### 3.2 20 years prediction and virtual variety testing recommendations

The architecture of the PTW model allowed to treat images as optional input. Hence, we could predict cultivar performance based on genetic marker data and historical weather data for a 20-year period across 10 locations in Switzerland. The PTW model demonstrated moderate alignment of average per-year yield predictions and measured yield reported by the Swiss interbranch organization Swiss granum (Figure 3D). However, certain years deviated from this trend: in 2016, the model predicted moderate yields despite exceptionally low observed values. The year 2016 was characterized by low-light conditions in Spring and an in general very wet season with low yields in all Europe (Le Gouis, Oury, and Charmet 2020), resulting in significant *Zymoseptoria tritici* pressure and resulting Septoria tritici blotch (STB) disease. Diseases are factors that the proposed PTW model can only incorporate indirectly via environmental covariates, which may lead to overestimations in situations with high disease pressure such as in 2016. In contrast, in 2011, 2014, and 2015, measured yields were notably higher than the model’s moderate predictions, for which there is no obvious cause known to us.

PTW model predictions were not only year-specific, but also location and genotype specific. Hence, the model allowed to virtually test varieties at unseen locations and years using historical weather data. The result of such a ‘virtual variety testing’ closely matched the official list of recommended varieties in 2018 published by Agroscope (Courvoisier et al. 2017) across all locations (Figure 3E). However, a closer examination at the individual location level revealed notable shifts in genotype rankings. For instance, while the variety ‘SIALA’ (a sister-line of CH CLARO) ranked among the top three on both the official list and the PTW model’s overall recommendation, its ranking varied significantly by location. For the location ‘Delley’, ‘SIALA’ was entirely outperformed by predictions for the variety ‘ARNOLD’, highlighting the potential for location-specific variety recommendations. However, the variety ‘ARNOLD’ also serves as an instructive example of how trial data may not always fully represent field-scale performance: Despite its initial promise on the recommended variety list in 2018, the variety was later removed due to its susceptibility to lodging under farming conditions.

### 3.3 P-state and G*×*E-state predictions

The PTW model extracts growth estimates from images (P-state) and from genotype-environment interactions based on temperature, VPD, and genetic marker data (G*×*E-state). Hence, the trained PTW can visualize those states and responses at inference, enabling an analysis and interpretation of learned genotype-characteristics.

The P-state encoder is based on images only and hence genetic marker data-agnostic, allowing to examine the genotype-specificity of the extracted P-state with a heritability calculation. Heritability is a common measure in plant breeding to indicate the extent to which the variation in a phenotypic trait has a genetic origin, with values close to one indicating a high dominance of the phenotypic signal (Falconer and Mackay 1996). The P-states exhibited varying heritability over time, with peaks coinciding with phenological stages of emergence, tillering, heading, and senescence (Figure 4A). P-state values demonstrated wide ranges throughout the season. Similarly, G*×*E-states varied, revealing clear genotype-specific differences (Figure 4B).

**Figure 4:**
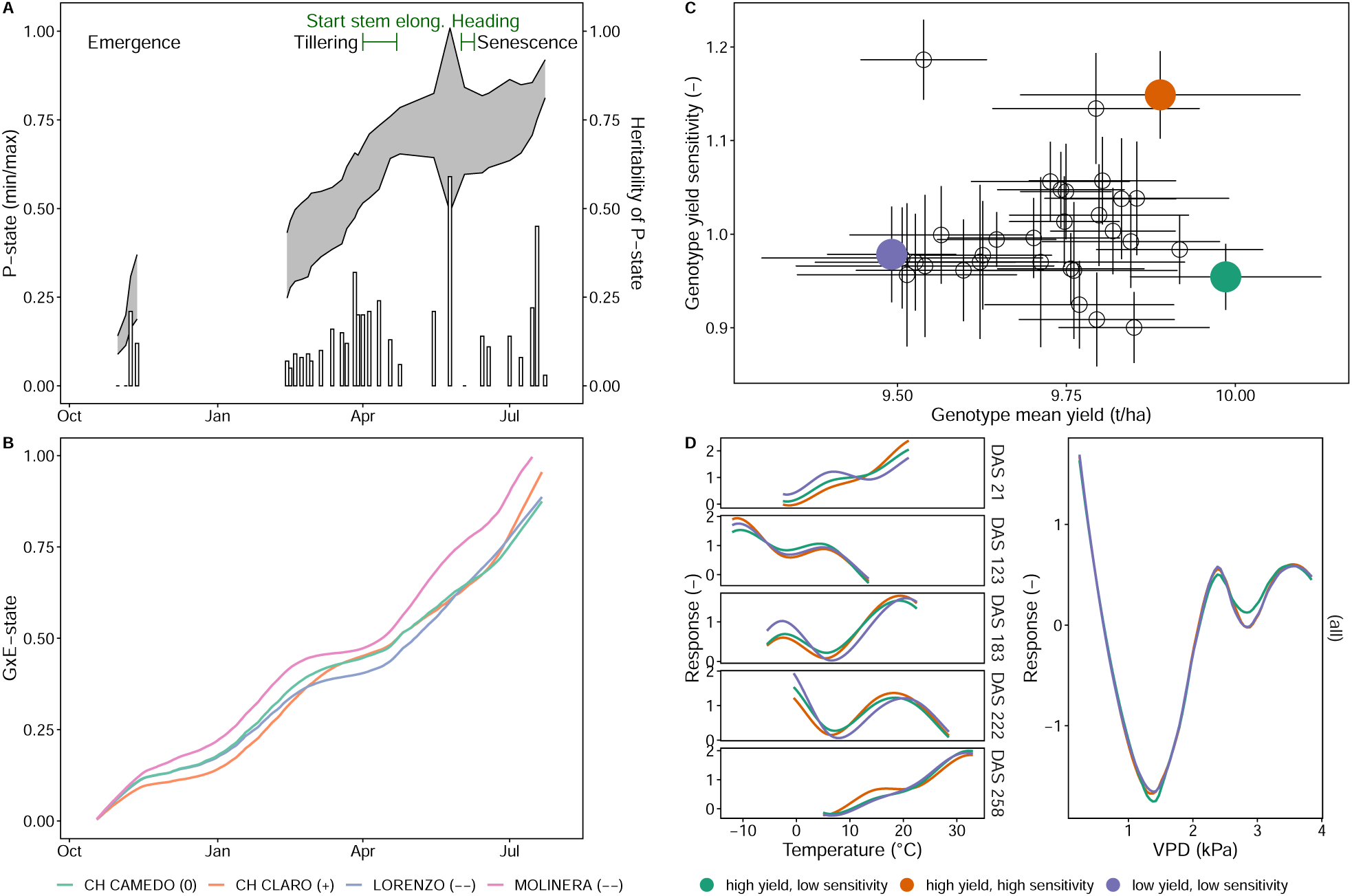
Predicted P-state and heritability for all varieties (A) and G*×*E-state of four selected winter wheat varieties sown in mid-October (B), and clustering of learned genotype responses (Splines) to temperature and vapour pressure deficit (VPD) in relation to their yield and yield sensitivity (C) for three exemplary clusters (D).

The learned temperature responses indicated non-linear responses to temperature with low growth rates at temperatures around 0 *^◦^*C, pronounced maximum growth rates at an optimum temperature around 20 *^◦^*C, and, depending on the growth phase, decreasing growth responses with supraoptimal temperatures above 20 *^◦^*C (Figure 4D). This response pattern changed throughout the season, with a tendency to higher optimum temperatures later in the season. In Winter (around 120 days after sowing (DAS)), an inversed response pattern emerged with higher responses at negative temperatures and lower responses at positive temperatures. Stress responses to VPD were characterised by an optimum VPD around 1.5 kPa and increased stress responses at lower and higher VPD values.

The fact that the GABI-MET data was unseen in training allowed to use this MET data to investigate genotype yield and genotype yield sensitivity using a Finlay-Wilkinson regression. Clustering of the learned genotype-specific responses to temperature and VPD resulted in groups that distinctly separated in this yield – yield sensitivity space (Figure 4C). Three clusters were visually sampled at the extremes of this space, one with high yield and low sensitivity, one with high yield and high sensitivity, and one with both low yield and low sensitivity. The clusters displayed distinguished response patterns to temperature and VPD (Figure 4D).

Genotypes in the high yield/low yield sensitivity group—the cluster that breeding is supposedly most interested in—were associated with reduced stress response to VPD around 1.5 kPa. Both genotype groups with high yield (insensitive and sensitive ones) were characterized by pronounced temperature responsiveness at emergence (DAS 21) and in Spring (DAS 183–222). Yet, during senescence, temperature responses of the high-yielding sensitive genotype cluster was more pronounced for temperatures below 20 *^◦^*C and less pronounces above 20 *^◦^*C than for the insensitive cluster.

## 4 Discussion

High-throughput field phenotyping platforms and multi-environment trials (METs) represent distinct data sources with complementary strengths: phenotyping platforms provide high temporal resolution (Kirchgessner et al. 2017), while METs offer high spatial resolution across geographical locations. METs can sample crucial combinations of climatic and soil conditions, but typically face a large *p*, small *n* problem. Field phenotyping platforms can deliver large data sets, but struggles with out-of-domain issues. Merging these two data sources has the potential to offset each other’s limitations, thereby improving predictions for unseen environments.

Yet, integrating multi-modal data such as images, genetic marker data, and environmental covariates is non-trivial. Our proposed PTW model achieves this integration by learning specific data encoders, with results demonstrating superior performance for unseen MET environments if compare to a state-of-the-art baseline.

Limited interpretability is one of the strongest limitations of most deep learning models. In contrast to this prevailing trend, the proposed PTW model allows visualization of growth responses related to temperature and VPD stress, hence providing insights in learned plant physiology mechanistics. The observed patterns aligned well with existing domain knowledge. Increasing stress due to VPD was identified starting from 2 kPa, and temperature responses showed seasonal variations in cardinal temperatures, which strongly agrees with previous research (Schoppach and Sadok 2013; Porter and Gawith 1999). The observed “inverted” temperature response in winter can be interpreted as vernalization requirements (Slafer 1996).

Overall, it should be noted that the extracted temperature response patterns indicate strong shifts associated with changing growth processes. While the base temperature of growth shifts from zero degrees Celsius at emergence to close to ten degrees Celsius at heading, optimum temperatures are pronounced during the early generative stage but appear negligible in the vegetative phase and towards senescence. Although not at the core of this work, one may still speculate based on these results that gene expression underlying temperature response characteristics shifts strongly over time.

It was often assumed that evolutionary processes have limited variations in response patterns within species (Parent and Tardieu 2012). Yet, the PTW model identified clear genotype-specific deviations from the well-known general response patterns. Those patterns were not only related to individual genotypes, but also to clusters of high-yielding genotypes with low respectively high sensitivity to environment differences. For a given target population of environments (TPE) where one can determine yield sensitivity beforehand, the model allowed providing new insights into optimized ideotypes. For the investigated TPE in this work (Switzerland), a selection for more responsive temperature reactions at emergence and during the senescence phase may lead to higher yield. Yield stability might be enhanced by selecting for higher resistance to VPD around 1.5 kPa.

In addition to assisting in ideotype formulation, a deep learning model such as the PTW may also directly support selection decisions in breeding. The model’s ability to genomically predict yield performance could enhance breeders’ decisions when selecting corresponding genotypes in their breeding programs through a genomic selection approach.

However, for any application it is crucial to ensure that the TPE is well-represented in the training data. Training solely on the FIP 1.0 data set and predicting “distant” environments in Germany and France showed significant bias and reduced agreement on genotype ranking. Adding MET data from an unrelated TPE did not improve results. This strongly indicates the difficulty of learning stress factors that are underrepresented in the training environment. We assume that combining multiple field phenotyping platform data sets that cover different stressors and growth conditions could substantially increase the power of such models. With the FIP 1.0 data set, one such data set is publicly available, and hopefully more will follow soon. Given recent developments in mobile and/or portable device phenotyping, breeders may have tools at hand to extend such data sets to better cover their TPE.

A striking finding of this study is the comparable performance of the baseline (random regression genomic prediction approach) with the PTW model when trained and tested on the FIP 1.0 data set, which comprises only one location. For such a scenario, genetic and environmental main effects are the main contributing components to yield, while G*×*E interaction effects are small (Roth et al. 2025). While the linear mixed model framework used for the random regression model is, in its core, perfectly suited for such a decomposition of measured variances into the components G, E, and G*×*E, we have to acknowledge that our current model architecture disentangles genotypic effects (G) from genotype-by-environment interactions (G*×*E), but does not explicitly separate environmental effects (E) from G*×*E. The G*×*E-state encoder lacks an explicit mechanism to structure the latent space into distinct components corresponding to G, E, and their interaction. A clearer separation of these three contributions could be achieved through alternative approaches, such as for example an autoencoder-style backbone in which phenomic images from the same genotype across multiple environments are grouped and jointly encoded to yield structured latent representations at multiple levels: general features shared across all images, environment-specific features common to similar conditions, and image-specific features capturing local variability. Such end-to-end representation learning strategies that explicitly integrate the disentanglement of G, E, and G*×*E effects may represent an important direction for follow-up work.

From a crop growth modelling point of view, the question arises how ‘unique’ individual model runs are in regard to learned plant physiological responses to environmental covariates, but also in regard to predictions to unseen environments. In other words, do the inherent characteristics of deep learning, equifinality and stochasticity, result in significantly different extracted responses? On the example of the two FIP 1.0 data set models trained on different training sets, this can not be observed—the extracted response curves per genotype are very comparable (see Supplementary Materials). Hence, we can conclude that while each learned model is unique, the learned responses to environmental covariant and predictions into unseen environments generalizes well.

Variety testing data have previously demonstrated the potential for location-specific variety recommendations (Herrera et al. 2018). However, variation across years in such data is usually significant and the reason why variety testing typically spans at least three years in most countries. Our work demonstrates how this time span can be further extended to “virtually” test in arbitrary historical years via a learned model. This approach enables fine-grained genotype recommendations based on the frequency a variety results as a top-ranked genotype per year-location. Given the availability of genetic marker data, such a model could be incrementally extended each year, effectively leveraging the substantial work produced by variety testing.

Traditional variety testing involves experiments across numerous locations, yet recommendations are typically based on overall adjusted genotype mean values to ensure statistical robustness. The proposed PTW model offers promising potential to enhance these recommendations by making them location-specific, potentially increasing crop production sustainability through locally adapted agricultural practices. By integrating MET data with location-specific descriptive factors such as management practices and soil types, the PTW model could effectively distinguish between relatively stable location effects and variable environmental factors that change year to year. This separation would likely improve prediction accuracy for location-specific recommendations, providing more tailored guidance to producers.

Looking beyond the present, the proposed PTW model has the potential to enable predictions related to climate change for future climate scenarios. We refrained from pursuing this in our work. One reason for this decision was that projected temperature trajectories for future climate scenarios may exceed the conditions in the training data, making predictions unreliable. Such out-of-domain effects were observed in the results for German and French environments in general, and for specific years for the Swiss environment. The year 2016 for example with very low reported yield was characterized by low-light conditions in Spring and an in general very wet season in all Europe (Le Gouis, Oury, and Charmet 2020), conditions that may not be present in the FIP 1.0 data set used in training. Having field phenotyping data from environments that correspond to such condition could significantly help to address this limitation.

Another reason for not projecting results onto future climate scenarios was that the PTW model relies on hourly temperature and VPD values. Climate scenario projections on an hourly basis are still rare; for example, those from MeteoSwiss are currently available only on a daily basis (Fischer et al. 2022). However, this situation is expected to improve in the near-future.

In future works, predictions could be extended beyond yield to include quality traits such as grain protein content, Zeleny sedimentation values, and other relevant parameters. Such extensions would, of course, be dependent upon the availability of corresponding phenotypic data, a limitation of the data sets used in this work. Yet, the end-to-end deep learning approach exemplified by the PTW model could be readily adapted for multi-trait prediction with minimal additional complexity.

We conclude that the integration of high-throughput field phenotyping platform data with multi-environment trial data in deep learning models represents a significant advancement in crop performance prediction. Our PTW model successfully bridged these complementary data sources, enabling more accurate predictions across diverse environments, and revealing important genotype-specific responses to environmental stressors. The proposed approach offers valuable insights for developing optimized crop ideotypes and enhancing breeding decisions. While challenges remain in extrapolating to environments beyond the ones learned from the training data, the framework established here provides a foundation for future work, including potential applications in climate change adaptation strategies. As more comprehensive data sets become available and climate projections improve in resolution, the predictive power and applicability of such integrated models will continue to expand, ultimately contributing to more resilient and productive agricultural systems.

## Data availability

Data from the field phenotyping platform 1.0 (FIP 1.0) platform have been released as the FIP1.0 data set, https://doi.org/10.1093/gigascience/giaf051. The GABI-MET data set is public, https://doi.org/10.5447/ipk/2022/18. The CH-MET data set was at the time of writing subject to limitations that may have been resolved in the meantime. Please contact Agroscope (Juan Herrera) for further details.

## Code availability

All code are available from the gitlab repository of ETH Zurich—Model code repository: https://gitlab.ethz.ch/crop_phenotyping/PTW_model; Data preparation code repository: https://gitlab.ethz.ch/crop_phenotyping/PTW_annotate. Model weights of trained models are provided with the code respository https://gitlab.ethz.ch/crop_phenotyping/PTW_model (‘./scripts/weights’). Preprocessed input batches and all model weights are available upon request (> 1.5 TB).

## Supporting information

Supplementary materias

## Abbreviations Acronyms

R1: auto-regression of order 1
ASReml-R: statistical package that fits linear mixed models using Residual Maximum Like- lihood (REML).
BLUE: best linear unbiased estimation
CDC: Climate Data Center
CH-MET: Swiss variety testing multi-environment trial
DAS: days after sow-ing
DNA: deoxyribonucleic acid
DWD: Deutscher Wetterdienst
FIP: field phenotyping platform
FIP 1.0: field phenotyping platform 1.0
GABI-WHEAT: Genomanalyse im Biologischen System Pflanze - Weizen
GABI-MET: GABI multi-environment trial
MAPE: mean absolute percentage error
MET: multi-environment trial
MeteoSwiss: Federal Office of Meteorology and Climatology MeteoSwiss.
MSE: mean squared error
PTW: plant time warping
RGB: red, green, and blue
SNP: single-nucleotide polymorphism
SPEI: standardized precipitation and evapotranspiration index
SYNOP: surface synoptic observation
TPE: target population of environments
VPD: vapour pressure deficit.

## Acknowledgements

The authors would like to thank the Crop Science Group of ETH Zurich, especially Achim Walter, Andreas Hund and Norbert Kirchgessner, for their massive support in carrying out the field trials and phenotyping experiments, and Delley Semences et Plants SA (DSP), especially Flavio Foiada, for their essential support in preparing seed batches for the Swiss varieties.

## Authors contributions

L.R.: Conceptualization, Methodology, Software, Validation, Formal analysis, Investigation, Data Curation, Visualization, Funding acquisition, Investigation, Writing – original draft. J.M.H.: Methodology, Validation, Resources, Data Curation, Writing - Review & Editing. L.L.H.: Resources, Data Curation, Writing - Review & Editing. D.P.: Project administration, Resources, Writing - Review & Editing. D.F.: Project administration, Resources, Writing - Review & Editing. M.B.: Methodology, Writing - Review & Editing X.C.: Methodology, Writing - Review & Editing P.N.: Methodology, Writing - Review & Editing M.V.: Methodology, Resources, Supervision, Project administration, Writing - Review & Editing

## Extended data

### Extended tables

**Table 2:** Ablation study of PTW model, context FIP 1.0 test set, GABI SNP set.

| Model | Pearson’s r (correlation) |  |  | MAPE |  |  |
| --- | --- | --- | --- | --- | --- | --- |
|  | (E) | (G and E) | (G) | (E) | (G and E) | (G) |
| PTW | 0.32 | -0.08 | 0.35 | 31.5 | 24.8 | 9.4 |
| no L1 | 0.47 | 0.26 | 0.40 | 30.7 | 27.7 | 9.1 |
| no L2 | 0.00 | 0.00 | 0.00 | 12.7 | 11.7 | 17.9 |
| no L3 | 0.00 | 0.00 | 0.00 | 12.7 | 11.7 | 17.9 |
| no L4 | -0.01 | -0.30 | -0.13 | 41.6 | 39.6 | 26.6 |
| no P-state transformer | 0.45 | -0.34 | 0.38 | 16.3 | 12.6 | 11.3 |
| no main G effect | 0.14 | -0.06 | 0.13 | 18.0 | 16.8 | 9.9 |

### Extended figures

The following figures visualize predictions versus observed values for the three data sets (1) field phenotyping platform 1.0 (FIP 1.0) (Figure 5), (2) GABI multi-environment trial (GABI-MET) (Figure 6) and (3) Swiss variety testing multi-environment trial (CH-MET) (Figure 7).

**Figure 5:**
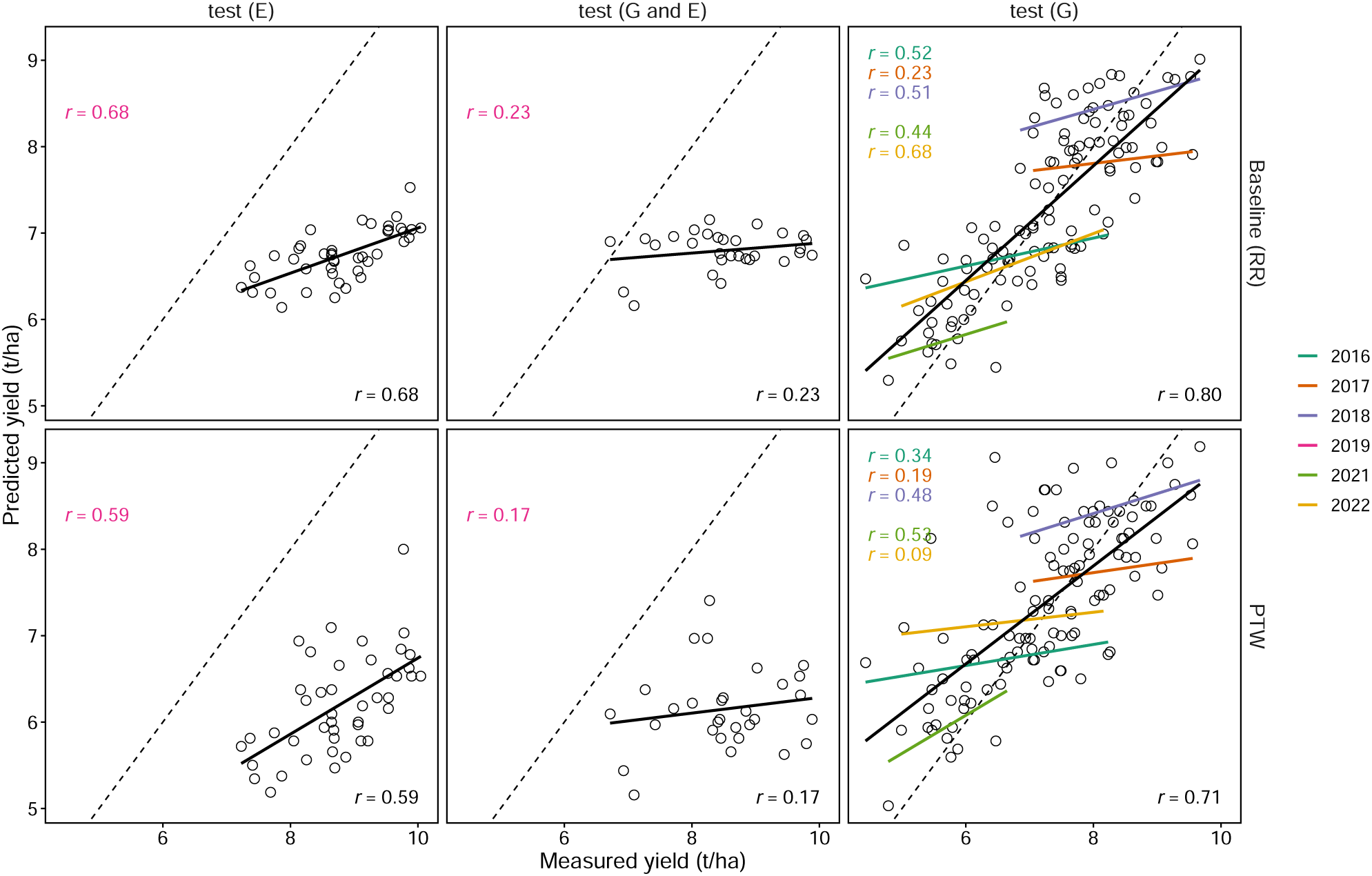
Predicted versus measured yield for the field phenotyping platform 1.0 (FIP 1.0) (Extended SNP marker) data set for the baseline (random regression, RR) and the plant time warping (PTW) model. The dashed line indicates the 1:1 reference.

**Figure 6:**
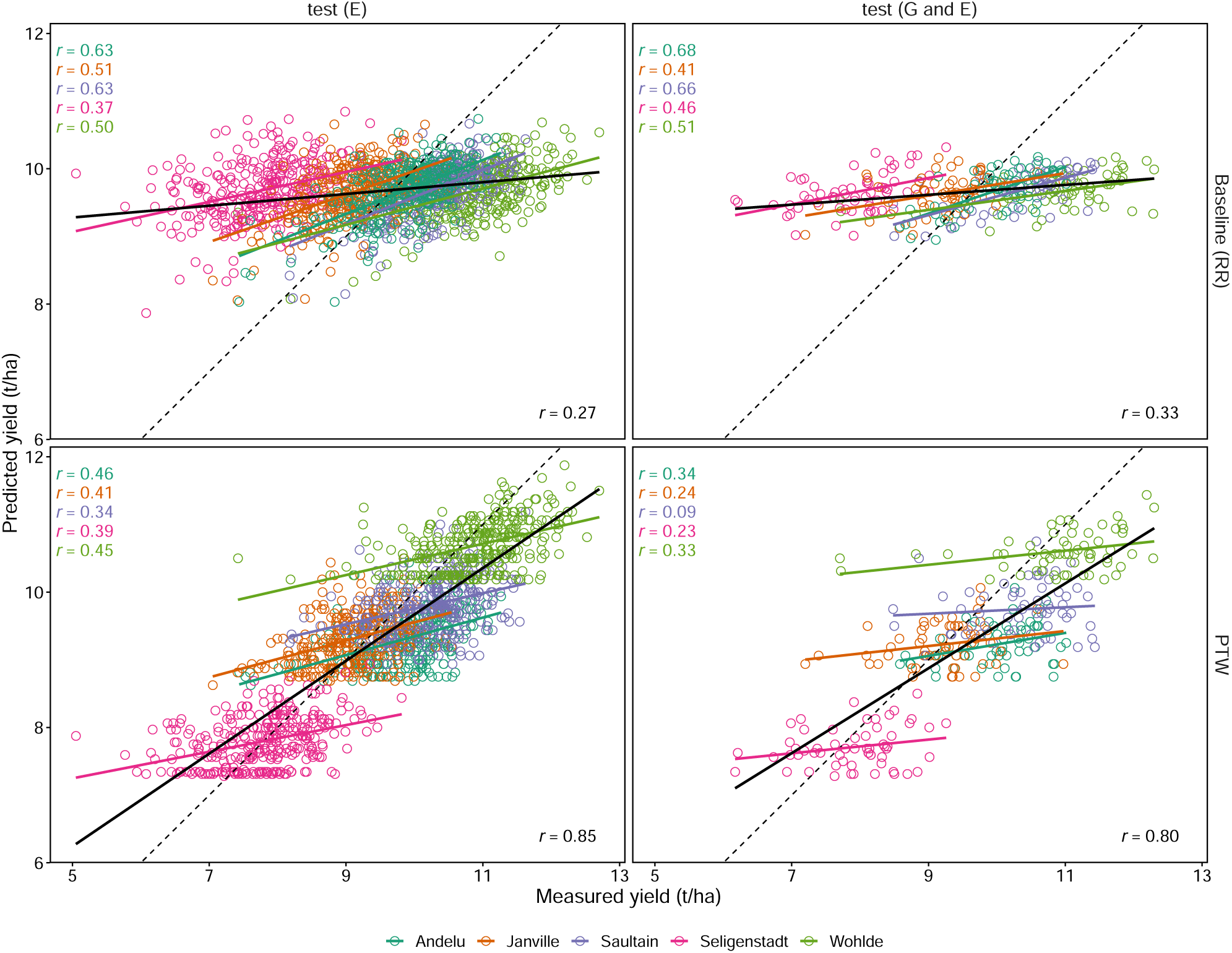
Predicted versus measured yield for the GABI multi-environment trial (GABI-MET) (2010) data set for the baseline (random regression, RR) and the plant time warping (PTW) model. The dashed line indicates the 1:1 reference.

**Figure 7:**
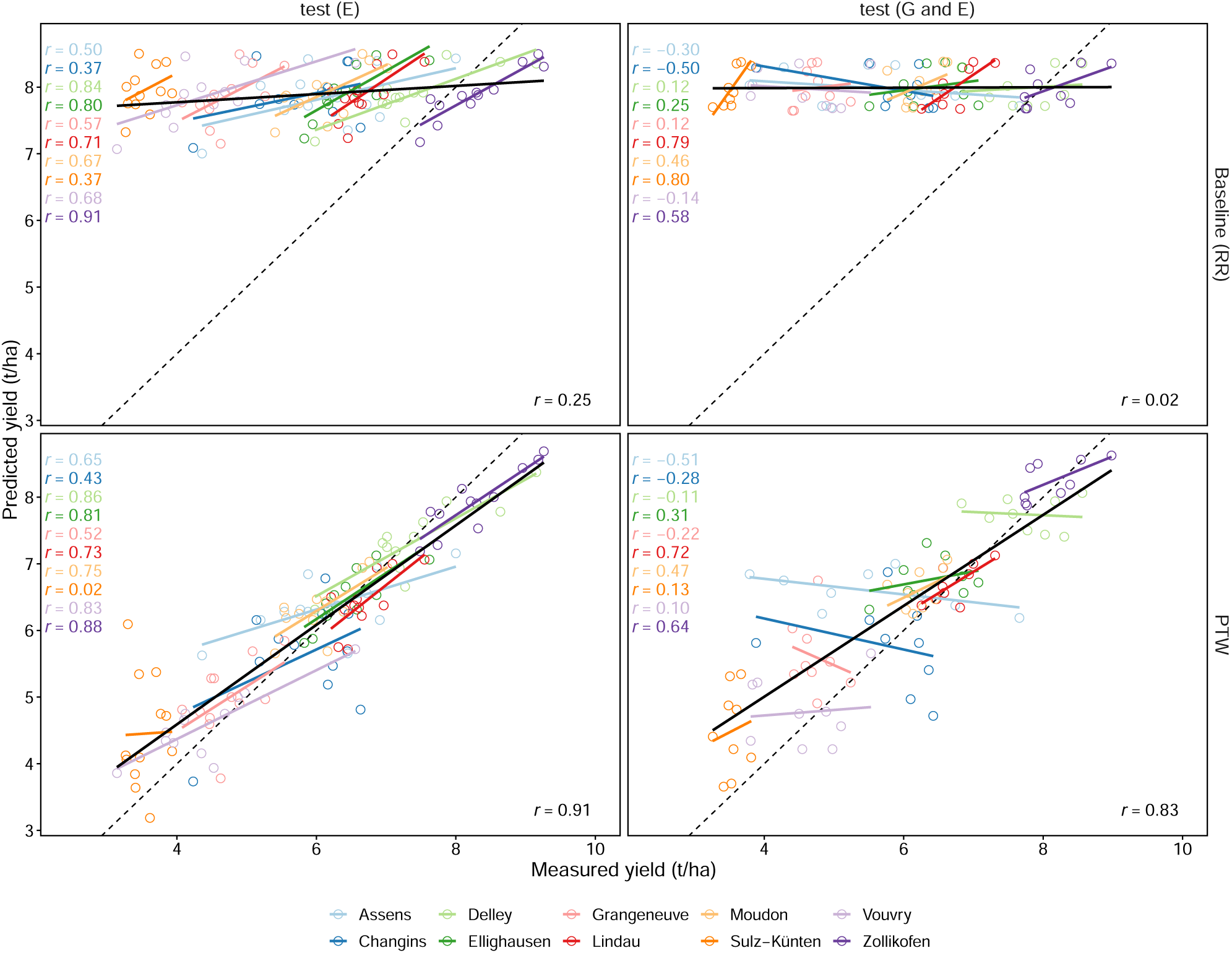
Predicted versus measured yield for the Swiss variety testing multi-environment trial (CH-MET) data (2013) set for the baseline (random regression, RR) and the plant time warping (PTW) model. The dashed line indicates the 1:1 reference.

