## Supplementary materias for "Characterizing yield through wheat’s perception of chronological progression: a multi-omics plant-time warping approach"

**Figure A1**

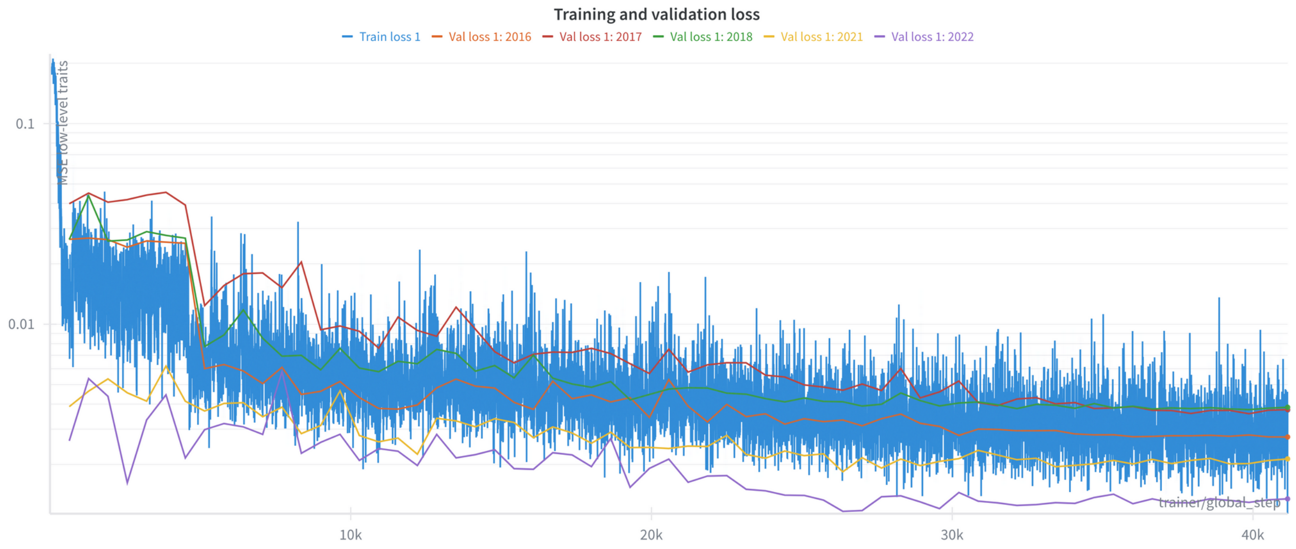


Training versus validation loss L1 (low-level traits) for model GABI-WHEAT SNP. Note that this is training step 1 (low-level trait training) before freezing low-level trait prediction for second training step.

**Figure A2**

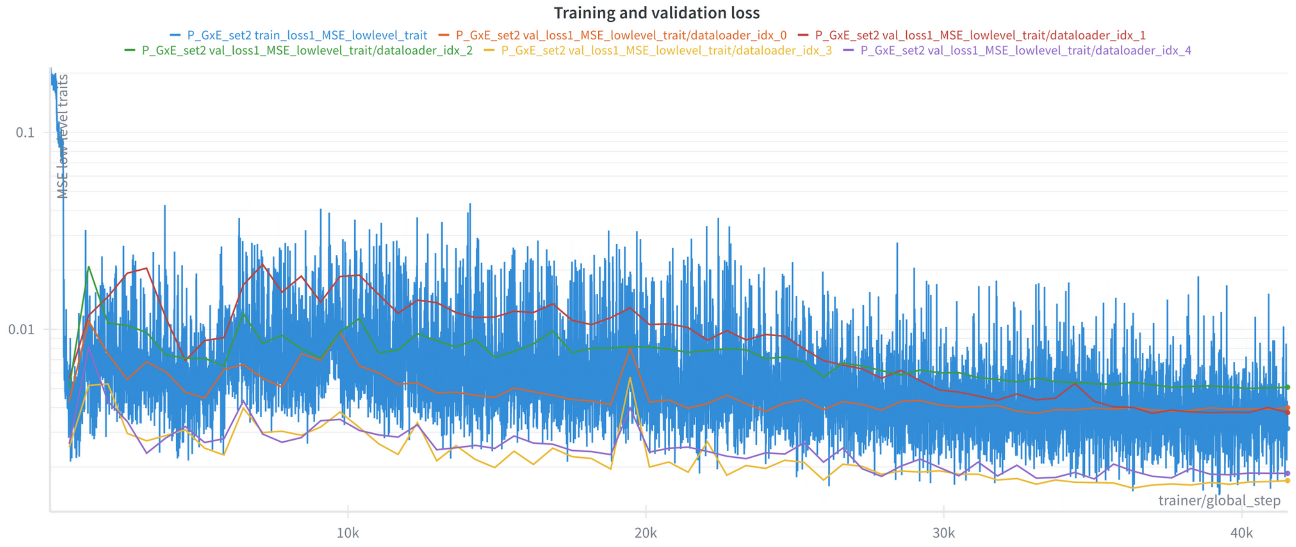


Training versus validation loss L1 (low-level traits) for model Extended SNP. Note that this is training step 1 (low-level trait training) before freezing low-level trait prediction for second training step.

### Figure A3

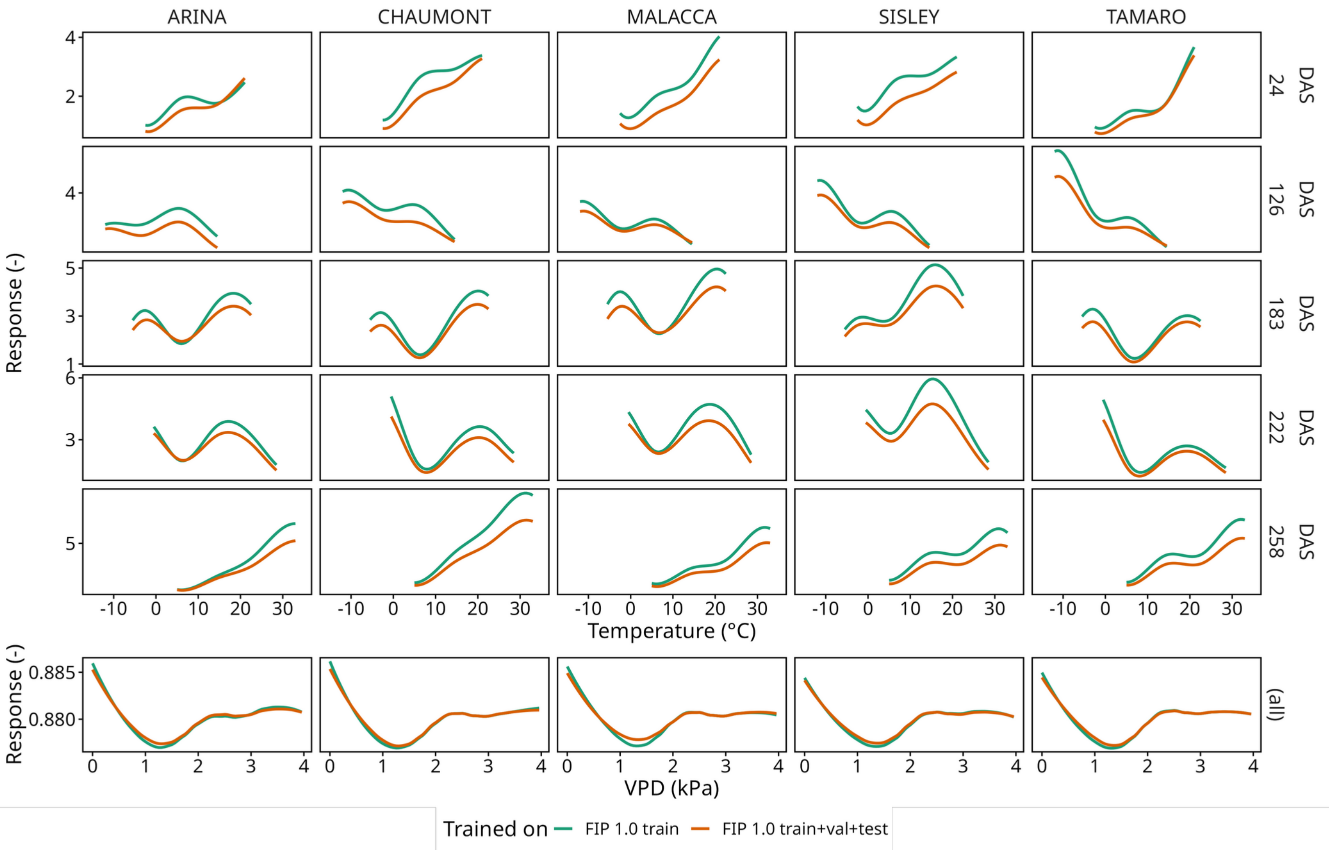


Learned response pattern to temperature and VPD for 5 randomly selected genotypes for two independently trained models. Model 1 (FIP 1.0 train) was trained on a subset of the data used for model 2 (FIP 1.0 train+val+test), initialization of model parameters was random.
